# Phospholipase C β4 promotes RANKL-dependent osteoclastogenesis by interacting with MKK3 and p38 MAPK

**DOI:** 10.1101/2024.03.19.585823

**Authors:** Dong-Kyo Lee, Xian Jin, Poo-Reum Choi, Ying Cui, Xiangguo Che, Sihoon Lee, Keun Hur, Hyun-Ju Kim, Je-Yong Choi

## Abstract

Phospholipase C beta (PLCβ) exerts diverse biological processes, including inflammatory responses and neurogenesis; however, its role in bone cell function is largely unknown. Among the PLCβ isoforms (β1–β4), we found that PLCβ4 was most highly upregulated during osteoclastogenesis. In this study, we used global knockout and osteoclast lineage-specific PLCβ4 conditional knockout (*LysM-PLCβ4^−/−^*) mice and demonstrated that PLCβ4 is a crucial regulator of receptor activator of nuclear factor κB ligand (RANKL)-induced osteoclast differentiation. Deletion of PLCβ4, both globally and in the osteoclast lineage, resulted in a significant reduction in osteoclast formation and the downregulation of osteoclast marker genes. Importantly, *LysM-PLCβ4^−/−^* male mice exhibited greater bone mass and a lower number of osteoclasts *in vivo* than their wild-type littermates, without altering osteoblast function. Mechanistically, we found that PLCβ4 forms a complex with p38 mitogen-activated protein kinase (MAPK) and MAPK kinase 3 (MKK3) in response to RANKL, thereby modulating p38 activation. An immunofluorescence assay further confirmed the colocalization of PLCβ4 with p38 after RANKL exposure. Moreover, p38 activation rescued the impaired osteoclast formation and restored the reduced p38 phosphorylation due to *PLCβ4* deficiency. Thus, our findings reveal that PLCβ4 controls osteoclastogenesis via the RANKL-dependent MKK3-p38 MAPK pathway, and PLCβ4 may be a potential therapeutic candidate for bone diseases such as osteoporosis.

## Introduction

Osteoclasts, which originate from monocyte/macrophage lineage cells, are multinucleated cells that are primarily responsible for bone resorption. The differentiation process of osteoclasts from their precursors requires two essential cytokines: macrophage colony-stimulating factor (M-CSF) and receptor activator of nuclear factor κB ligand (RANKL) (1–5). Osteoclastogenesis is initiated through the interaction of RANKL with its receptor RANK, which leads to the recruitment of the adaptor molecule TNF receptor-associated factor 6 (TRAF6). This RANK/TRAF6 complex triggers the activation of TGF-β-activated kinase-1 (TAK1), which in turn activates the IκB kinase (IKK) complex and mitogen-activated protein kinases (MAPKs), including p38, c-Jun N-terminal protein kinase (JNK), and extracellular signal-regulated kinase (ERK) (6, 7). The activation of these signaling cascades ultimately results in the expression and stimulation of nuclear factor of activated T cells cytoplasmic 1 (NFATc1), a key transcription factor for osteoclast differentiation (8). During the final stage of osteoclastogenesis, NFATc1 interacts with several transcription factors, including AP-1, PU.1, and microphthalmia transcription factor (MITF), to induce the expression of osteoclast-specific genes, such as tartrate-resistant acid phosphatase (TRAP) and cathepsin K (CTSK) (9).

Among the three MAPKs, p38 MAPK is a major kinase that plays an essential role in osteoclast differentiation and bone homeostasis. Mice with an osteoclast lineage-specific deletion of p38α that was generated through the LysM-Cre system exhibit increased bone mass, which is associated with a reduced number of osteoclasts and bone resorption (10). Similarly, mice with Mx-Cre-mediated p38α conditional knockout (KO) reveal increased bone mass because of the reduced number of osteoclasts and are protected against TNF-α-mediated inflammatory bone destruction (11). In addition, specific pharmacological p38 inhibitors have been shown to prevent inflammatory- or ovariectomy-induced bone loss in animal models (12–14). Expression of the dominant negative form of p38 or treatment with p38 inhibitors completely blocks RANKL-induced osteoclast formation from osteoclast precursor cells *in vitro*, further supporting these findings (15–17). Furthermore, p38 MAPK directly stimulates NFATc1 and MITF, thereby inducing the expression of osteoclastogenic markers (18, 19). These studies emphasize the critical role of p38 MAPK activation by RANKL as an essential signaling event required for osteoclast differentiation.

The phospholipase C (PLC) family functions as a critical regulator of phosphoinositide metabolism by hydrolyzing phosphatidylinositol-4,5-bisphosphate into inositol-1,4,5-triphosphate and diacylglycerol (20). These products function as essential secondary messengers that govern various cellular processes. Dysregulation of PLC function or expression can result in various disorders, such as cancer, neurological disorders, and immune dysfunction. The PLC family, which is comprised of 13 members, can be categorized into 6 subfamilies (β, γ, δ, ε, ζ, and η) based on their domain structure and mode of activation (21–24). Within the PLC family, the PLCβ subfamily is further comprised of four isozymes (β1, β2, β3, and β4). The expression levels of PLCβ isoforms vary according to tissue type. PLCβ2 is expressed exclusively in hematopoietic cells, whereas PLCβ1 and PLCβ3 are found in various cells and tissues. On the other hand, PLCβ4 is primarily expressed in neuronal and retinal tissue (25–27) and has a well-established role in neurological function. Mice that lack PLCβ4 exhibit central nervous system defects such as ataxia, absence seizures, and anxiety behavior (28–31). PLCβ4 deficiency also leads to defects in phototransduction and visual processing (32, 33). In addition, a recent study identified PLCβ4 as a critical regulator of TCR signaling and immune response (34), thus highlighting its importance in immune function. Furthermore, mutations in PLCβ4 are associated with a human disease known as auriculocondylar syndrome, also referred to as question mark ear syndrome, in which patients have developmental defects in their ears and mandible (35, 36). Despite these intriguing findings, the role of PLCβ4 in bone cells, particularly in osteoclasts, remains unknown.

This study found that PLCβ4 showed the highest level of upregulation among the PLCβ isoforms during osteoclast differentiation. Therefore, we investigated the role of PLCβ4 in RANKL-induced osteoclastogenesis and bone homeostasis by employing a gene knockdown technique and using global KO and osteoclast lineage-specific conditional KO mice for PLCβ4.

## Results

### PLCβ4 expression is upregulated during osteoclastogenesis

To explore the potential factors that affect osteoclastogenesis, bone marrow-derived macrophages (BMMs) were cultured with M-CSF and RANKL for 4 days to induce osteoclast generation (Fig. 1a). Following this, we performed global transcriptomic analysis using both BMMs and mature osteoclasts. The microarray analysis data revealed a strong induction of PLCβ4 by RANKL, as well as the upregulation of various osteoclastogenic genes (Fig. 1b). To verify these results, we examined the expression patterns of PLCβ isoforms by real-time PCR. The mRNA levels of PLCβ1 and PLCβ3 decreased during osteoclastogenesis, whereas the mRNA expression of PLCβ4 was most significantly upregulated as the cells differentiated (Fig. 1c). Consistently, the protein level of PLCβ4 was elevated throughout the process of osteoclastogenesis (Fig. 1d).

**Fig. 1.**
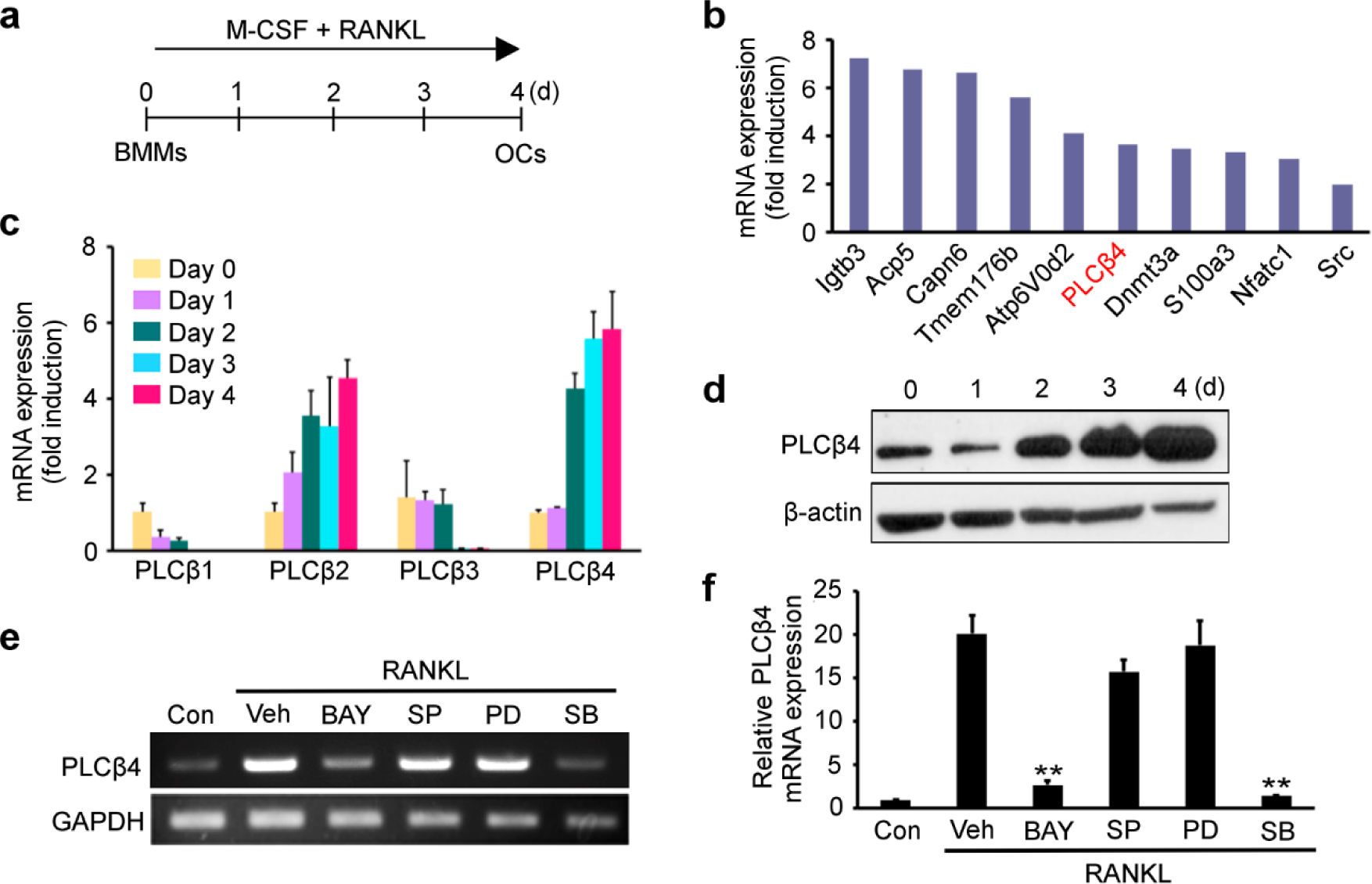
Expression of PLCβ4 is upregulated during osteoclastogenesis. **a** Bone marrow-derived macrophages (BMMs) were cultured with 30 ng/mL of macrophage-colony stimulating factor (M-CSF) and 20 ng/mL of receptor activator of nuclear factor κB ligand (RANKL) for 4 days to induce osteoclasts (OCs) differentiation. **b** Microarray profile of PLCβ4 expression during osteoclastogenesis. **c** Quantitative real-time PCR was performed to assess the mRNA expression levels of the PLCβ isoforms (β1–β4) during osteoclastogenesis. **d** PLCβ4 protein levels were determined by immunoblotting. **e**, **f** BMMs were cultured with M-CSF and RANKL in the presence of the indicated inhibitors. PLCβ4 expression levels were analyzed by reverse transcription PCR (**e**) or real-time PCR (**f**). BAY, Bay11-7802 (1 μM); SP, SP600125 (20 μM); PD, PD98059 (20 μM); SB, SB203580 (20 μM). Data are expressed as the mean ± standard deviation (SD). \*\**P* < 0.001 vs. the vehicle

We further determined which signaling cascades were involved in the upregulation of PLCβ4 expression by RANKL. To achieve this, BMMs were cultured with M-CSF and RANKL in the presence of various inhibitors of the RANKL signaling pathways, such as NF-κB and MAPKs. Our results showed that the RANKL-mediated upregulation of PLCβ4 expression was mainly via the NF-κB and p38 MAPK pathways, as the increased PLCβ4 expression was significantly abolished by the NF-κB inhibitor Bay11-7802 and the p38 inhibitor SB203580 (Fig. 1e, f). However, the JNK inhibitor SP600125 and the MEK inhibitor PD98059 did not affect the RANKL-mediated PLCβ4 upregulation (Fig. 1e, f).

### PLCβ4 knockdown reduces osteoclast differentiation

The upregulation of PLCβ4 expression via RANKL suggests its potential importance in osteoclasts. To explore this, we employed the lentiviral-mediated expression of a small hairpin RNA (shRNA) to silence PLCβ4 expression. As shown in Fig. 2a, the lentiviral-mediated PLCβ4-shRNA efficiently reduced the mRNA expression of PLCβ4 in BMMs (day 0) and preosteoclasts (day 2). The transduced BMMs were then cultured with M-CSF and two different concentrations of RANKL for 4 or 5 days. The knockdown of PLCβ4 significantly inhibited the formation of TRAP-positive multinucleated cells at both of the RANKL concentrations (Fig. 2b, c) and decreased TRAP activity in the culture medium (Fig. 2d). However, PLCβ4 knockdown did not affect the proliferation of osteoclast precursors, as determined by an MTS assay (Fig. 2e). We further assessed the impact of PLCβ4 knockdown on the expression of osteoclast-specific markers. PLCβ4 knockdown significantly attenuated the induction of various osteoclastogenic genes, such as NFATc1, TRAP, cathepsin K (CTSK), Atp6V0d2, and OSCAR (Fig. 2f).

**Fig. 2.**
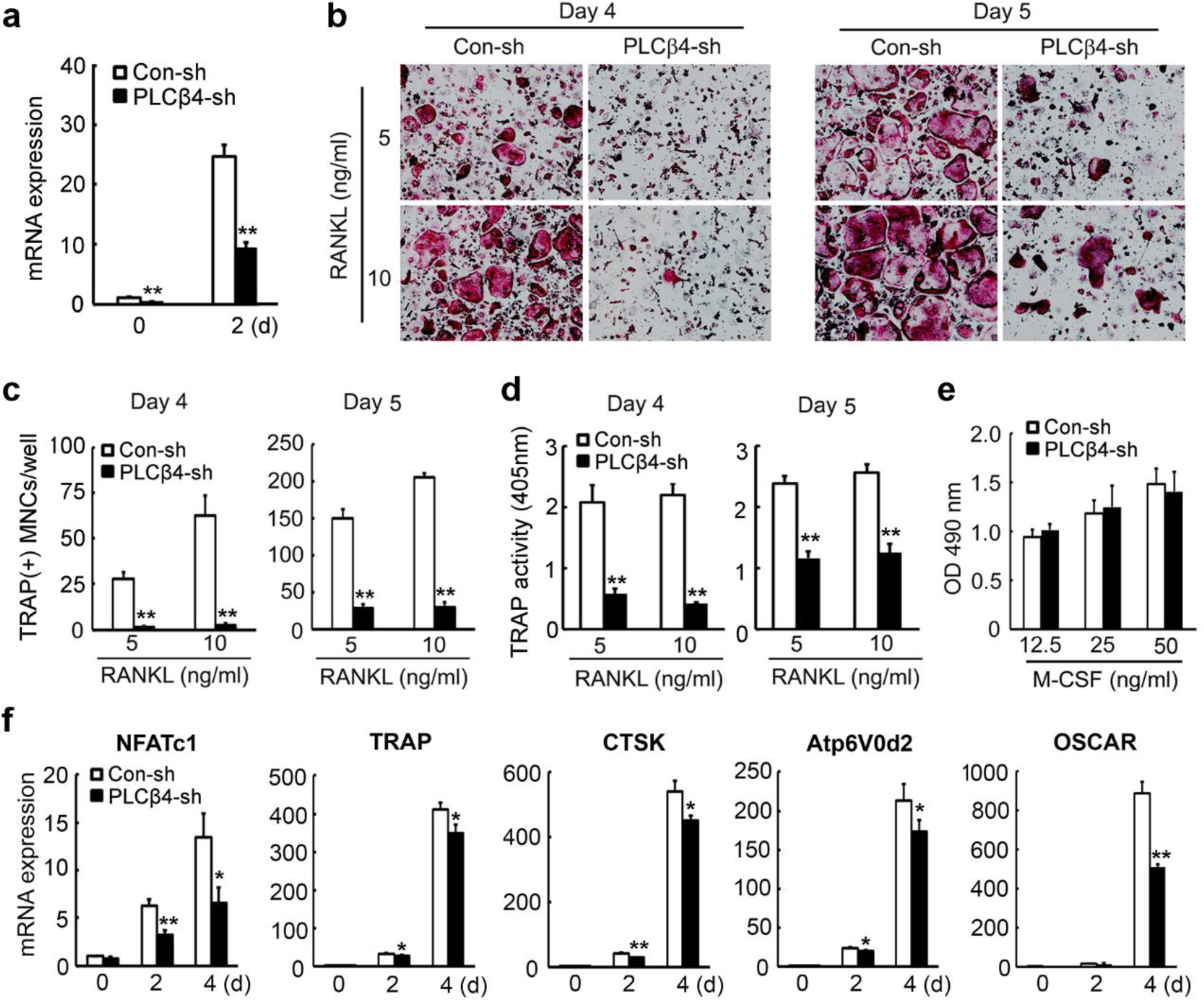
PLCβ4 knockdown reduces RANKL-mediated osteoclast differentiation. BMMs were transduced with non-specific control shRNA (Con-sh) or PLCβ4 shRNA (PLCβ4-sh) lentiviral particles. **a** The expression of PLCβ4 was analyzed by real-time PCR in BMMs (day 0) or preosteoclasts (day 2). **b**, **c**, **d** Transduced BMMs were cultured with M-CSF (30 ng/mL) and the indicated doses of RANKL for 4 or 5 days. **b** Osteoclasts were visualized after tartrate-resistant acid phosphatase (TRAP) staining and, **c** the number of osteoclasts was counted. **d** TRAP activity was determined by using the cultured cell supernatant generated in (**b)**. **e** Transduced BMMs were cultured in the presence of various concentrations of M-CSF. After 3 days, the MTS assay was performed, as described in the Materials and Methods section. **f** Transduced BMMs were cultured in osteoclastogenic media for the indicated times. Real-time PCR was performed to assess the gene expression of osteoclastogenic markers. All data are expressed as the mean ± SD. \**P* < 0.05 and \*\**P* < 0.001 vs. the control (Con-sh)

### Global deletion of *PLCβ4* decreases osteoclastogenesis

Having established that PLCβ4 knockdown reduces osteoclastogenesis, we used PLCβ4 global KO (*PLCβ4^−/−^*) mice to further investigate the impact of PLCβ4 deficiency in osteoclasts. Interestingly, the size of 8-week-old *PLCβ4^−/−^* mice was smaller than their wild-type (WT) counterparts (Fig. 3a). However, except for mandibular development, no notable differences in skeletal development were observed between the WT and *PLCβ4^−/−^* mice at embryonic day 17.5, as determined by alcian blue and alizarin red staining (Fig. 3b). With regard to osteoclastogeneis, we first confirmed the PLCβ4 deletion in the *PLCβ4^−/−^*BMMs through immunoblotting analysis (Fig. 3c). Subsequently, we cultured BMMs from WT and *PLCβ4^−/−^* mice in osteoclastogenic media for 4 or 5 days and then performed TRAP staining. The results showed a significant reduction in the number of osteoclasts in *PLCβ4^−/−^* mice compared with their WT littermates (Fig. 3d, e). Consistently, the gene expression of osteoclast markers, including NFATc1, TRAP, OSCAR, DC-STAMP, and Atp6V0d2, was downregulated in the *PLCβ4^−/−^* cells compared with the WT cells (Fig. 3g). However, there was no significant difference in the MITF expression. We further observed that the loss of *PLCβ4* does not influence the proliferation of osteoclast precursors (Fig. 3f).

**Fig. 3.**
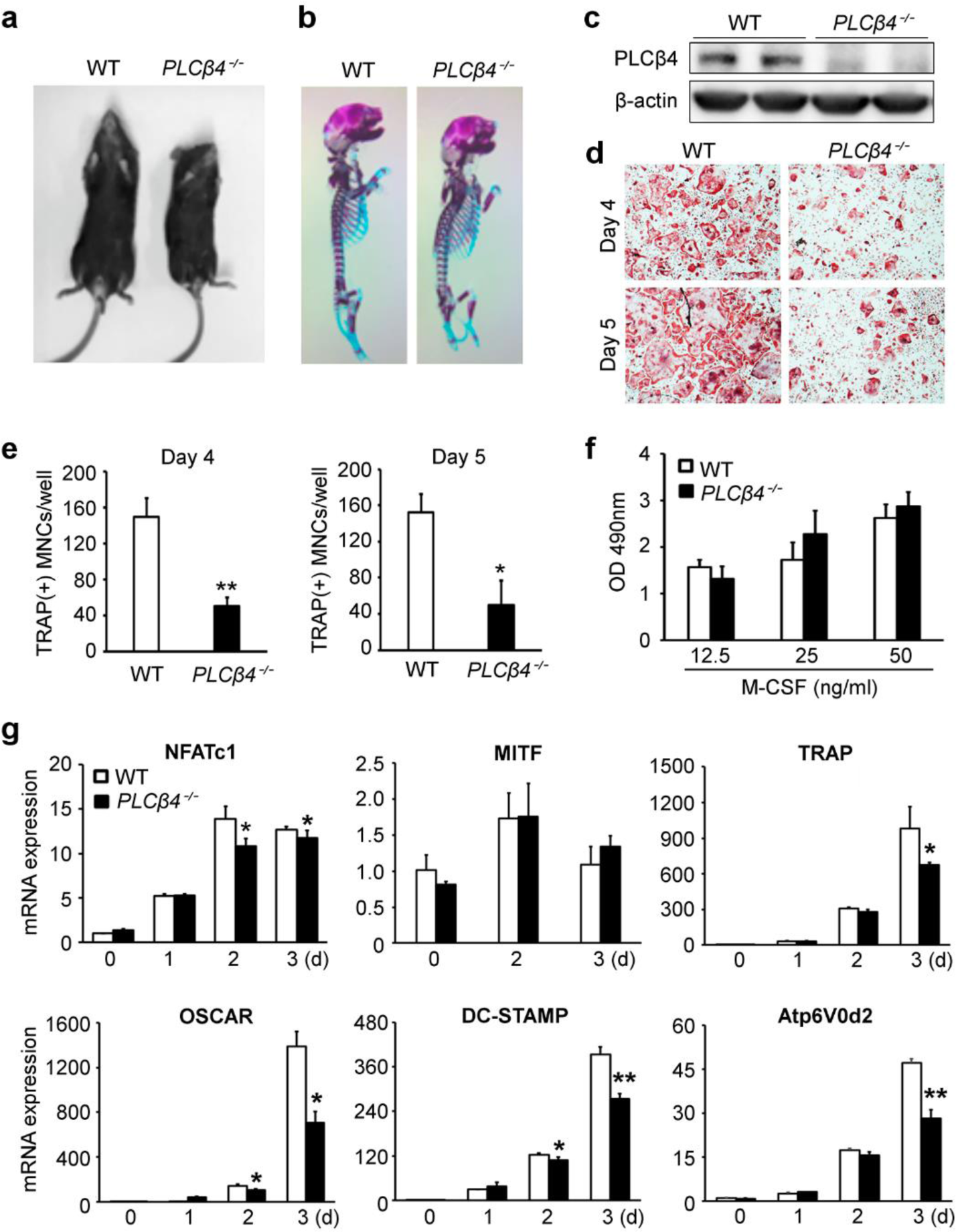
Global deletion of *PLCβ4* reduces osteoclastogenesis. **a** Size comparison of 8-week-old wild-type (WT) and *PLCβ4^−/−^*mice. **b** Alcian blue and alizarin red staining of the skeleton of WT and *PLCβ4^−/−^* mice at embryonic day 17.5. **c** Deletion of PLCβ4 in *PLCβ4^−/−^* BMMs was confirmed by immunoblotting. **d**, **e** The BMMs from WT and *PLCβ4^−/−^* mice were cultured with M-CSF (30 ng/mL) and RANKL (20 ng/mL) for 4 or 5 days. **d** The cultured cells were fixed and subsequently stained for TRAP. **e** Quantification of the number of osteoclasts. **f** An MTS assay was performed, and the absorbance was measured at 490 nm. **g** The BMMs from WT and *PLCβ4^−/−^* mice were cultured in the presence of M-CSF and RANKL for the indicated days. The expression of osteoclast marker genes was measured by real-time PCR. All data are expressed as the mean ± SD. \**P* < 0.05 and \*\**P* < 0.001 vs. the WT

### Osteoclast lineage-specific deletion of *PLCβ4* reduces osteoclastogenesis

To better understand PLCβ4 involvement in osteoclastogenesis, we generated osteoclast lineage-specific PLCβ4 conditional KO mice. To achieve this, we crossed *PLCβ4* floxed mice with LysM-Cre mice, which resulted in *LysM-Cre;PLCβ4^f/f^* mice (hereafter referred to as *LysM-PLCβ4^−/−^*). For comparison, we used littermates that were homozygous for the *PLCβ4^f/f^* genes but lacked the LysM-Cre allele, and this group was designated as the control (Control). The deletion of PLCβ4 in BMMs and osteoclasts obtained from *LysM-PLCβ4^−/−^* mice was confirmed by immunoblotting (Fig. 4a). In addition, we observed that both male and female *LysM-PLCβ4^−/−^* mice did not exhibit any significant gross morphological changes and had similar body weights as their control littermates (Fig. 4b).

**Fig. 4.**
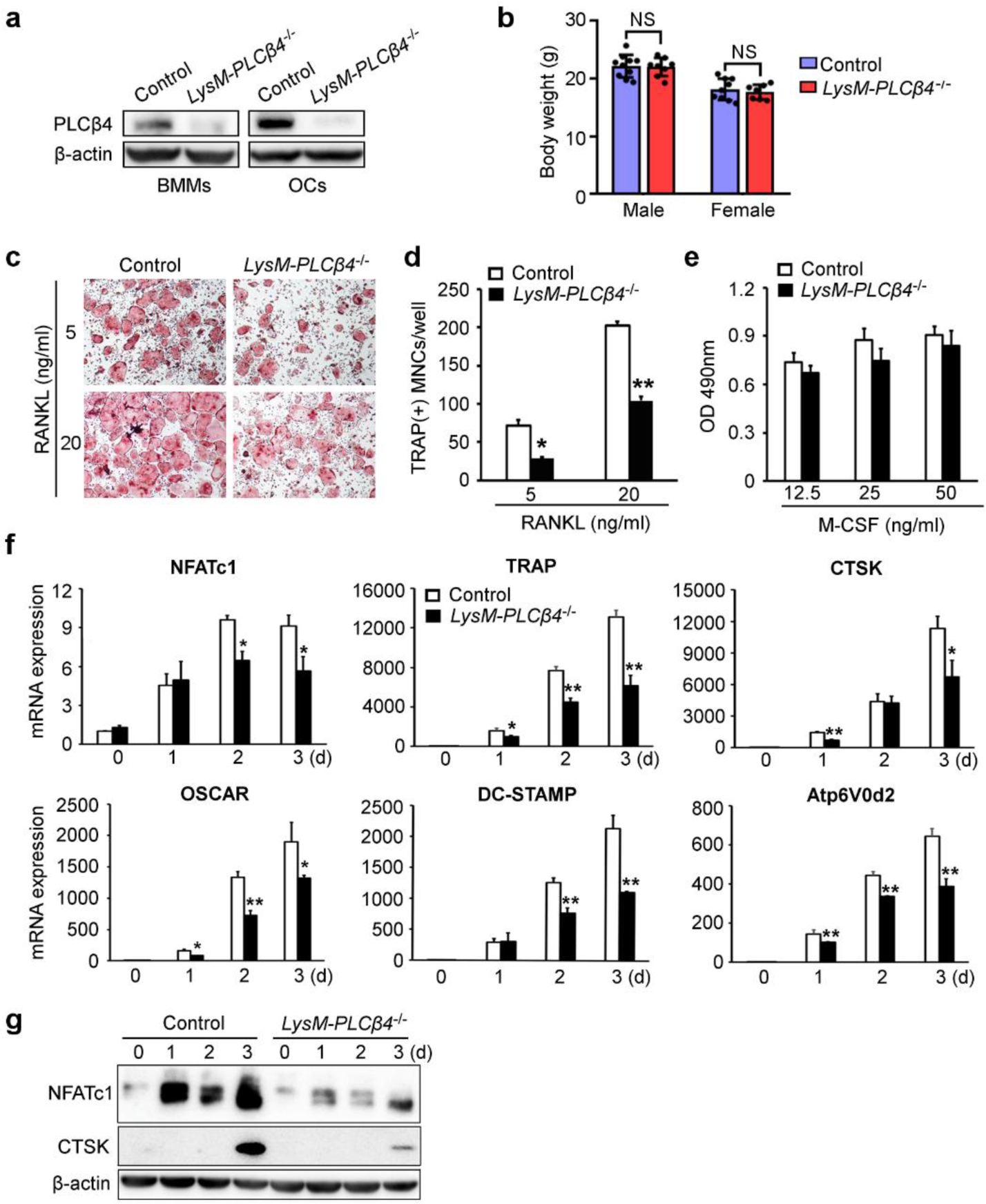
Conditional deletion of *PLCβ4* in the osteoclast lineage reduces osteoclastogenesis. **a** Deletion of PLCβ4 in *LysM-PLCβ4^−/−^* BMMs and osteoclasts (OCs) was confirmed by immunoblotting. **b** Comparison of the body weight of 8-week-old control and *LysM-PLCβ4^−/−^*male and female mice (n = 7–10). **c**, **d** The BMMs from the control and *LysM-PLCβ4^−/−^* mice were cultured with the indicated concentrations of RANKL in the presence of M-CSF (30 ng/mL). **c** Osteoclasts were visualized after TRAP staining. **d** Quantification of the number of osteoclasts. **e** An MTS assay was performed, and the absorbance was measured at 490 nm. **f**, **g** BMMs from the control and *LysM-PLCβ4^−/−^* mice were cultured in osteoclastogenic media for the indicated days. The expression of the indicated genes was measured by real-time PCR (**f**) or immunoblotting (**g**). All data are expressed as the mean ± SD. \**P* < 0.05 and \*\**P* < 0.001 vs. the control; NS, not significant

To assess the impact of osteoclast lineage-specific PLCβ4 deficiency on osteoclastogenesis, we cultured BMMs from the control and *LysM-PLCβ4^−/−^* mice in the presence of M-CSF and two different concentrations of RANKL. At both concentrations of RANKL, the number of osteoclasts was significantly reduced in the *LysM-PLCβ4^−/−^* group compared with that of the control group (Fig. 4c, d). However, PLCβ4 deficiency did not alter osteoclast precursor proliferation (Fig. 4e). Consistent with the reduced formation of osteoclasts, in *LysM-PLCβ4^−/−^*cells there was a significant decrease in the mRNA expression of various osteoclast-specific marker genes, including NFATc1, TRAP, and CTSK, when compared to the control group (Fig. 4f). The reduced expression of osteoclast-specific genes was further supported by the marked reduction in the protein levels of NFATc1 and CTSK in the cells derived from *LysM-PLCβ4^−/−^* mice (Fig. 4g).

### *LysM-PLCβ4^−/−^* male mice exhibit increased bone mass and decreased osteoclast number

To investigate the role of PLCβ4 in bone homeostasis *in vivo*, we performed three-dimensional microstructural analysis using microcomputed tomography (μCT) on 8-week-old control and *LysM-PLCβ4^−/−^* male and female mice. Interestingly, male *LysM-PLCβ4^−/−^* mice exhibited a significant increase in trabecular bone mineral density (BMD), bone volume per tissue volume (BV/TV), and trabecular number (Tb.N). In contrast, no significant changes were observed in the female mice compared with their control littermates (Fig. 5a, b). This increase in bone parameters in male *LysM-PLCβ4^−/−^* mice was accompanied by a reduction in trabecular separation (Tb.Sp). However, there were no notable differences in the trabecular thickness (Tb.Th) (Fig. 5a, b). We further found that the deletion of PLCβ4, specifically in osteoclasts, had no impact on the cortical bone in either gender (Fig. S1). Increased bone mass in the male *LysM-PLCβ4^−/−^* mice was confirmed by non-decalcified bone histology. Von Kossa staining of lumbar vertebrae sections revealed a significant increase in BV/TV and Tb.N and a corresponding decrease in Th.Sp. In contrast, Tb.Th remained unchanged in *LysM-PLCβ4^−/−^* mice (Fig. 5c, d).

**Fig. 5.**
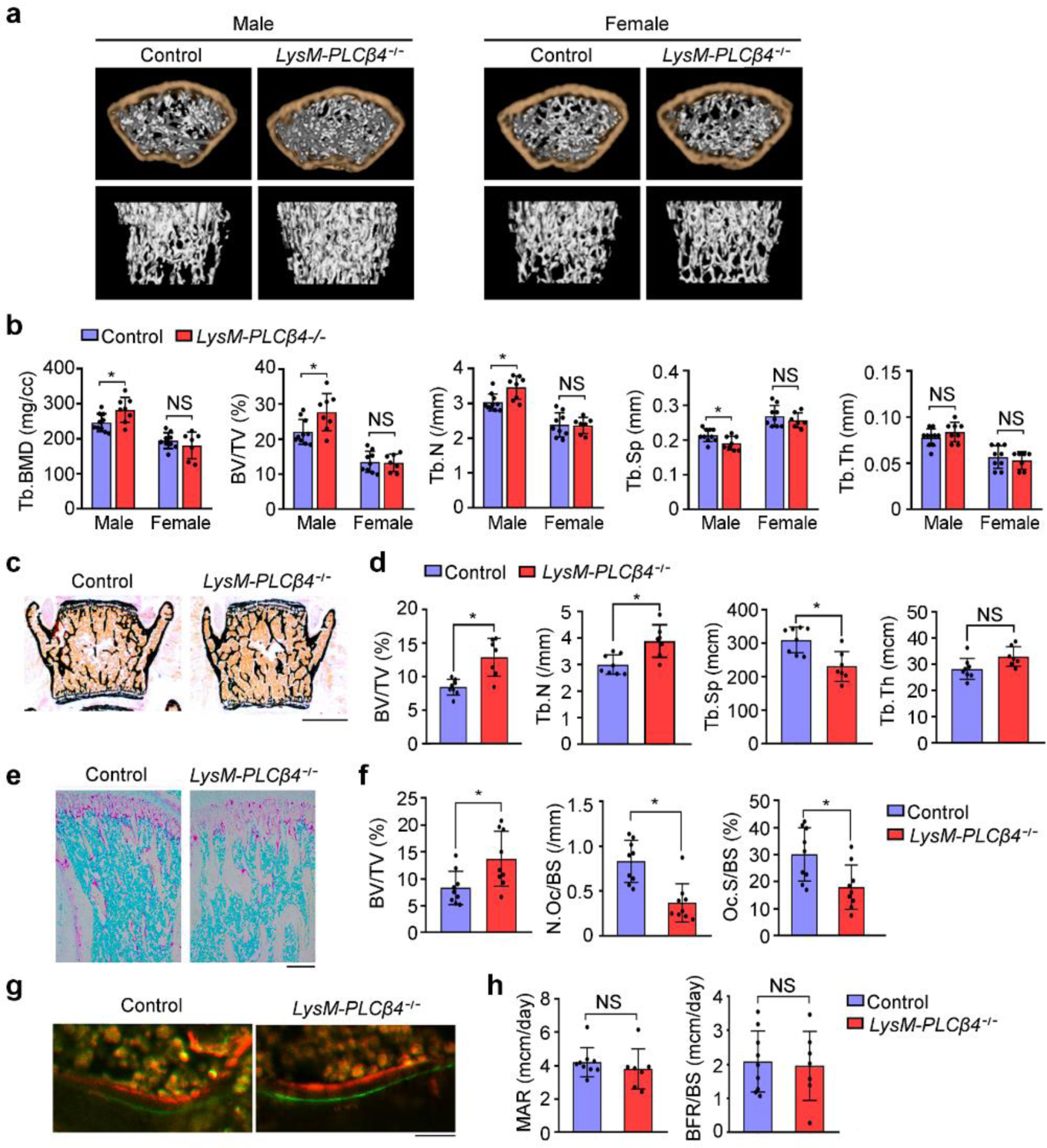
*LysM-PLCβ4^−/−^* male mice have increased bone mass. **a** Representative µCT images of femurs from 8-week-old control and *LysM-PLCβ4^−/−^* male and female mice. **b** Quantitative µCT analysis of trabecular bone parameters including bone mineral density (BMD), bone volume per tissue volume (BV/TV), trabecular number (Tb.N), trabecular separation (Tb.Sp), and trabecular thickness (Tb.Th) (n = 7–10). **c** Von Kossa staining of vertebral sections from 8-week-old control and *LysM-PLCβ4^−/−^*male mice. Scale bar = 1 mm. **d** Histomorphometric quantification of the data from (**c**) (n = 7–8). **e** TRAP staining images of the tibia from 8-week-old control and *LysM-PLCβ4^−/−^* mice. Scale bar = 200 μm. **f** Quantification of BV/TV, number of osteoclasts per bone surface (N.Oc/BS), and osteoclast surface per bone surface (Oc.S/BS) in (**e**) (n = 9). **g** Calcein and alizarin red double labeling images of trabecular bones in the vertebrae from 8-week-old control and *LysM-PLCβ4^−/−^* mice. Scale bar = 20 μm. **h** Mineral apposition rate (MAR) and bone formation rate (BFR/BS) were examined using histomorphometry of the images in (**g**) (n = 7–9). All data are expressed as the mean ± SD. \**P* < 0.05; NS, not significant

To examine whether the increased bone mass in *LysM-PLCβ4^−/−^*mice was due to the reduced number of osteoclasts, we performed histomorphometric analysis on the femurs of 8-week-old *LysM-PLCβ4^−/−^*male mice and their control littermates. Consistent with the μCT data, we found increased trabecular bone mass in *LysM-PLCβ4^−/−^* mice (Fig. 5e, f). TRAP staining revealed a significant reduction in the number of osteoclasts per bone surface (N.Oc/BS) and osteoclast surface per bone surface (Oc.S/BS) in *LysM-PLCβ4^−/−^* mice (Fig. 5e, f). However, when the bone formation parameters were examined using calcein and alizarin double labeling, we found no differences in the mineral apposition rate (MAR) and bone formation rate (BFR) between *LysM-PLCβ4^−/−^* mice and their control littermates (Fig. 5g, h). These results indicate that the targeted deletion of *PLCβ4,* specifically in the osteoclast lineage, leads to a decrease in osteoclastogenesis without altering osteoblast formation, which results in greater bone mass *in vivo*.

### PLCβ4 selectively modulates RANKL-mediated p38 MAPK

RANKL primarily controls osteoclastogenesis by activating downstream signaling pathways, including NF-κB and MAPKs (JNK, ERK, and p38). Therefore, we investigated the molecular mechanism by which PLCβ4 regulates osteoclastogenesis. Serum-starved BMMs from control and *LysM-PLCβ4^−/−^* mice were stimulated with RANKL. The phosphorylation of IκBα, JNK, and ERK in response to RANKL stimulation occurred normally in *LysM-PLCβ4^−/−^* BMMs (Fig. 6a, b). However, the activation of p38 was strongly blocked in *LysM-PLCβ4^−/−^* cells compared with the control BMMs (Fig. 6b). p38 MAPK mediates RANKL-induced osteoclast differentiation, partially through the phosphorylation of p65 at Ser-536, without affecting IκBα degradation (37). Thus, we further investigated whether PLCβ4 deficiency influenced p65 phosphorylation. We observed that RANKL-stimulated p65 phosphorylation at Ser-536 was significantly reduced in *LysM-PLCβ4^−/−^*BMMs (Fig. 6b). Because M-CSF and TNF-α also activate the p38 pathway, we next sought to determine whether these cytokines affect p38 signaling in PLCβ4-deficient cells. In contrast to RANKL stimulation, neither M-CSF nor TNF-α had an effect on p38 phosphorylation (Fig. 6c, d). These results demonstrate that PLCβ4 has a vital role in osteoclastogenesis by modulating the activation of the p38 signaling pathway in response to RANKL.

**Fig. 6.**
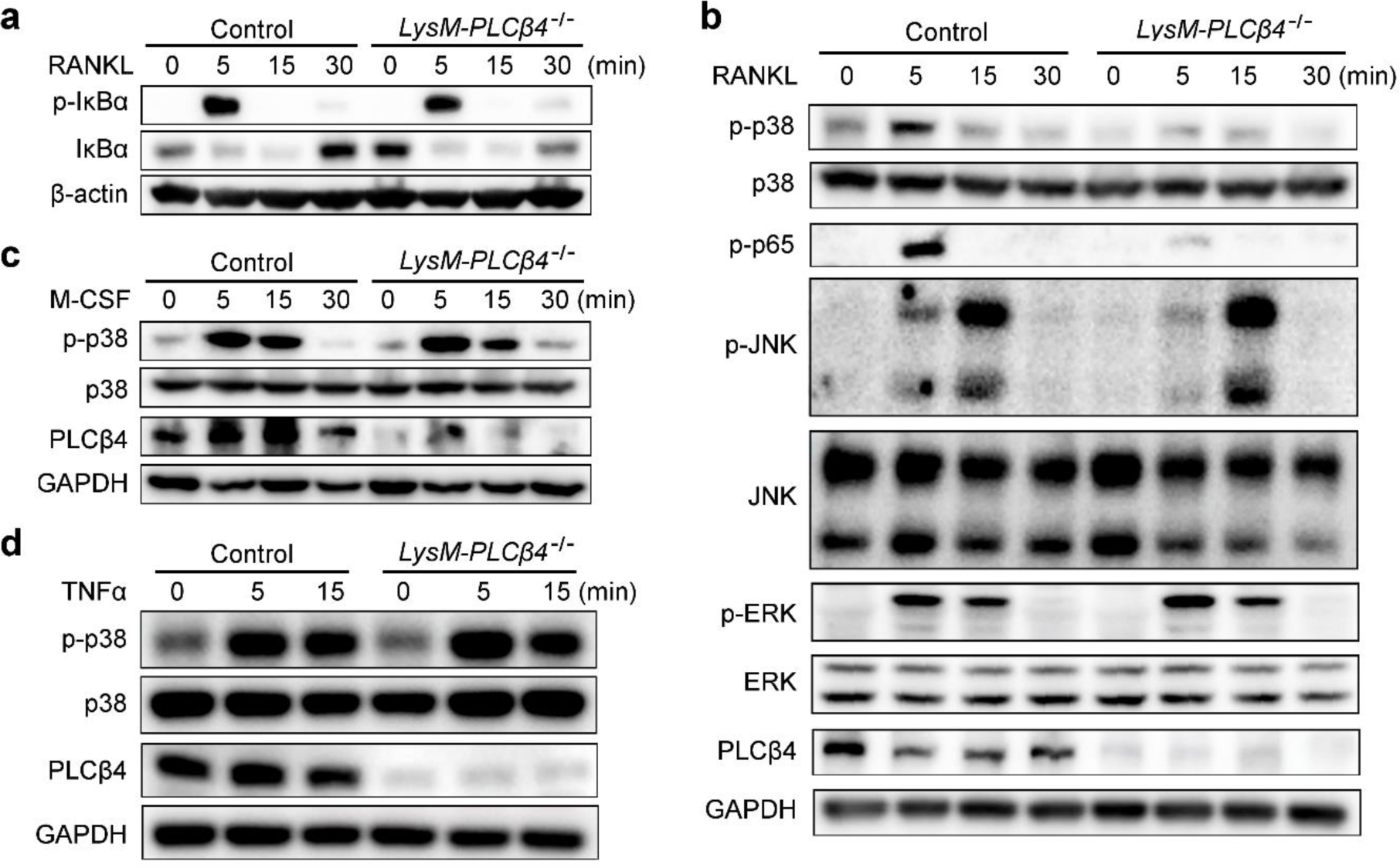
PLCβ4 modulates RANKL-mediated p38 phosphorylation. **a**, **b** Serum-starved BMMs from the control and *LysM-PLCβ4^−/−^* mice were stimulated with RANKL (50 ng/mL) for the indicated times. Immunoblotting was performed to assess IκBα phosphorylation and total IκBα (**a**) and the phosphorylation of p38, p65, JNK, and ERK (**b**). **c**, **d** Serum-starved BMMs from the control and *LysM-PLCβ4^−/−^* mice were stimulated with M-CSF (50 ng/mL) (**c**) or TNF-α (10 ng/mL) (**d**) for the indicated times and analyzed by immunoblotting with antibodies specific for the indicated proteins.

### PLCβ4 forms a complex with MAPK kinase 3 and p38 in response to RANKL

To investigate whether the catalytic activity of PLCβ4 is essential for RANKL-mediated p38 phosphorylation, control BMMs were pretreated with the PLC inhibitor U73122 and then exposed to RANKL stimulation. As shown in Fig. 7a, the PLC inhibitor did not affect p38 phosphorylation. This observation suggests that the catalytic activity of PLCβ4 is not required for p38 phosphorylation induced by RANKL. Interestingly, it has been reported that PLCβ interacts with p38 and MAPK kinase 3 (MKK3), an upstream regulator of p38, thus indicating the critical role of PLCβ as an interaction partner with MKK3 and p38 MAPK in cellular signaling processes (38). Based on these findings and our observation of reduced p38 phosphorylation in PLCβ4-deficient cells, we hypothesized that PLCβ4 can bind to p38 and modulate its phosphorylation. To explore this hypothesis, we conducted an immunoprecipitation assay using preosteoclasts and found that endogenous PLCβ4 associates directly with p38 (Fig. 7b). We then examined the impact of RANKL on the formation of the PLCβ4-MKK3-p38 complex. BMMs treated with RANKL were immunoprecipitated with an anti-PLCβ4 antibody. As shown in Fig. 7c, PLCβ4 was found to interact with MKK3 under basal conditions, and exposure to RANKL further enhanced this interaction, which led to the sequential recruitment of p38 to the PLCβ4-MKK3 complex. To confirm the interaction between PLCβ4 and p38 at the cellular level, we performed a cellular colocalization analysis of these two molecules in BMMs after RANKL stimulation. The results of immunofluorescence analysis revealed significant colocalization of PLCβ4 and p38 in RANKL-treated BMMs (Fig. 7d).

**Fig. 7.**
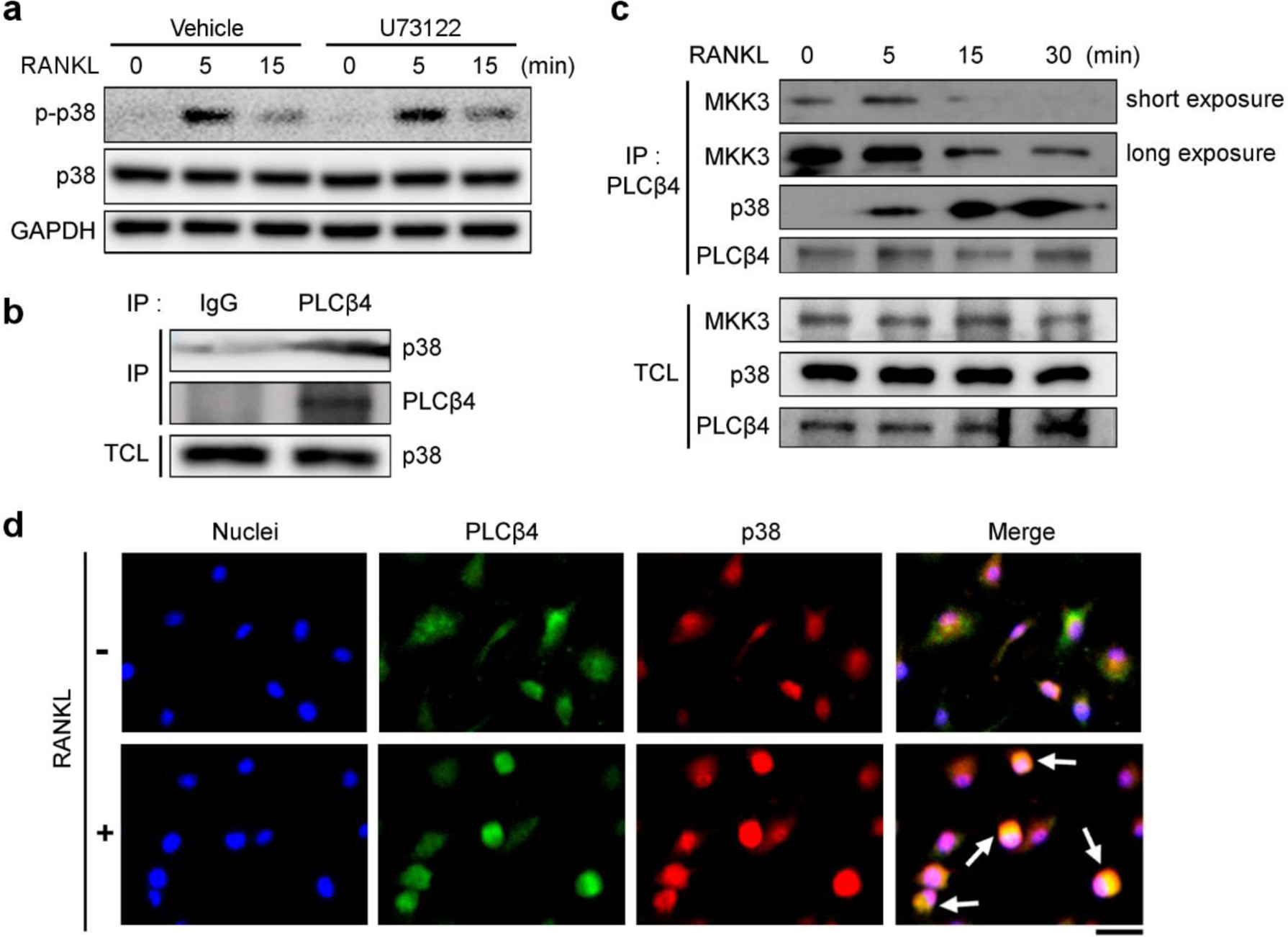
PLCβ4 interacts with p38 MAPK and MAPK kinase 3 (MKK3). **a** BMMs from the control and *LysM-PLCβ4^−/−^* mice were pretreated without or with the PLC inhibitor U73122 (5 μM) for 1 h and then stimulated with RANKL. Immunoblotting was performed to detect p38 phosphorylation in the cell lysates. **b** Endogenous PLCβ4 in preosteoclasts was immunoprecipitated and subjected to immunoblotting analysis using anti-p38 and anti-PLCβ4 antibodies. Non-specific proteins were immunoprecipitated with normal mouse IgG. **c** Serum- and cytokine-starved BMMs were stimulated with RANKL (50 ng/mL) for the indicated times and subjected to immunoprecipitation using an anti-PLCβ4 antibody, followed by immunoblotting for anti-MKK3, anti-p38, and anti-PLCβ4 antibodies. TCL, total cell lysate. **d** Serum- and cytokine-starved BMMs were stimulated without or with RANKL (50 ng/mL) for 15 min, fixed, and labeled with anti-PLCβ4 antibody (green), anti-p38 antibody (red), and Hoechst dye (blue). The images were detected using a fluorescence microscope. Scale bar = 20 μm.

### Activation of the p38 pathway rescues impaired osteoclastogenesis in PLCβ4 deficiency

Because PLCβ4 deficiency attenuates osteoclastogenesis by blocking p38 phosphorylation, we sought to determine whether the activation of the p38 pathway could rescue the impaired osteoclastogenesis observed in *LysM-PLCβ4^−/−^* mice. BMMs from control and *LysM-PLCβ4^−/−^* mice were pretreated with the vehicle or p38 activator anisomycin, followed by RANKL stimulation. The results revealed that the reduced p38 phosphorylation in *LysM-PLCβ4^−/−^*BMMs was restored at two different concentrations of the p38 activator (Fig. 8a). Moreover, treatment with a p38 activator rescued the decreased osteoclast formation in the absence of PLCβ4 (Fig. 8b, c).

**Fig. 8.**
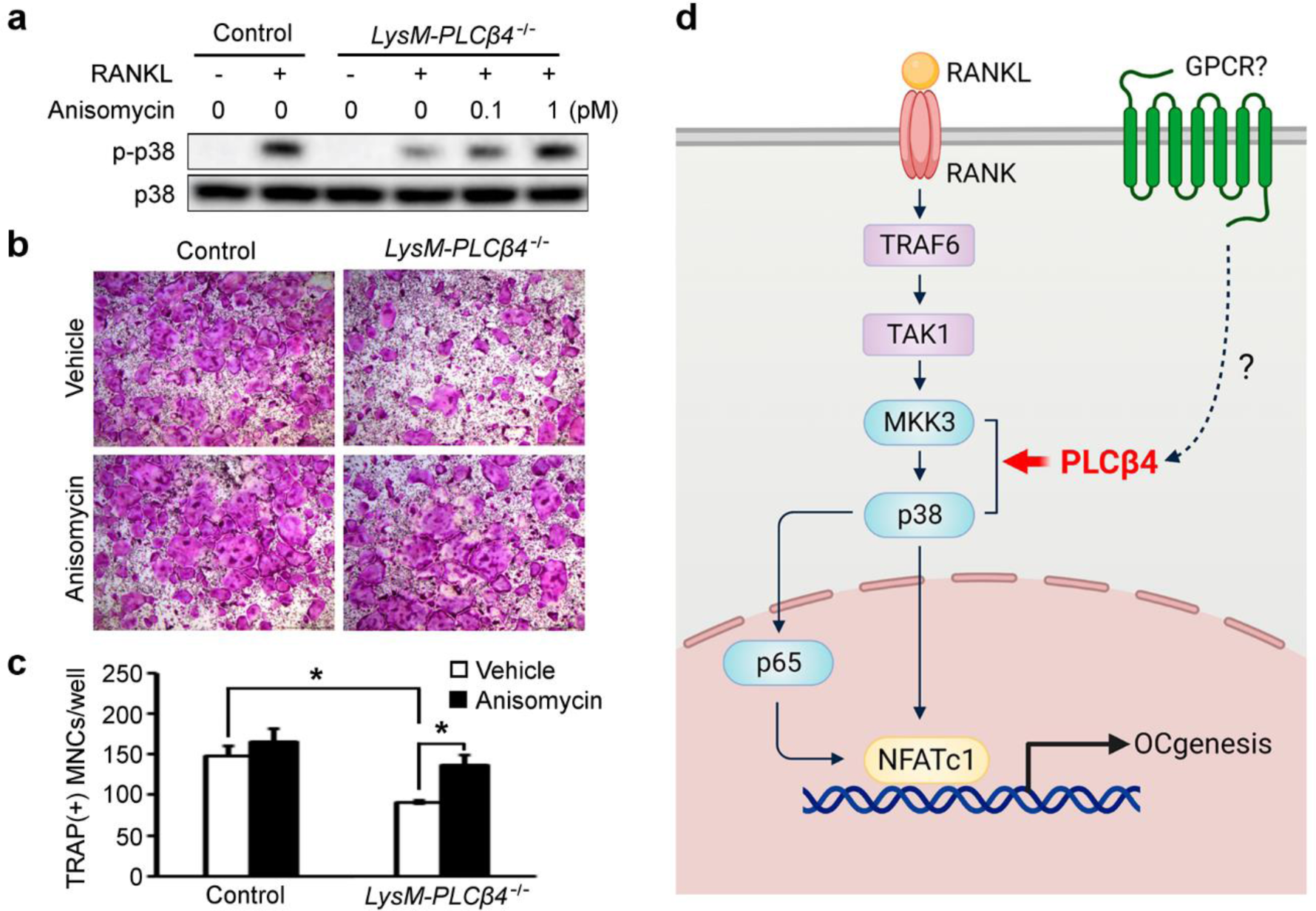
p38 activation restores attenuated osteoclast formation in *LysM-PLCβ4^−/−^* mice. **a** BMMs from the control and *LysM-PLCβ4^−/−^* mice were pretreated without or with the indicated concentrations of the p38 activator anisomycin for 2 h and then stimulated with RANKL (20 ng/mL) for 5 min. Immunoblotting was performed to detect p38 phosphorylation in the cell lysates. **b**, **c** BMMs from the control and *LysM-PLCβ4^−/−^* mice were cultured in osteoclastogenic media in the presence of anisomycin (1 pM). **b** Osteoclasts were visualized after TRAP staining. **c** Quantification of the number of osteoclasts. Data are expressed as the mean ± SD. \**P* < 0.05 vs. the vehicle. **d** Schematic diagram for the regulatory mechanism of PLCβ4 in osteoclastogenesis.

## Discussion

PLC is a key enzyme that maintains cellular homeostasis and normal physiological function. Among the PLC isozymes, PLCγ2 has been extensively studied and is recognized for its critical role in bone metabolism. The absence of PLCγ2 in mice leads to osteopetrosis because of defective osteoclastogenesis (39). The mechanism underlying this process involves PLCγ2 interacting with the immunoreceptor tyrosine-based activation motif (ITAM) and the adaptor molecule Gab2 to regulate osteoclast differentiation. Although the role of PLCγ2 in bone metabolism is well established, the functions of other PLC isozymes in bone homeostasis remain largely unexplored. In this study, we demonstrated that PLCβ4 plays a crucial role in regulating RANKL-mediated osteoclast differentiation. Our findings reveal that among the various PLC isozymes, PLCβ4, in addition to PLCγ2, is a critical regulator of bone homeostasis. Previous studies have demonstrated that PLCβ4 is prominently expressed in the brain and retina (25–27). A recent study also reported the detection of PLCβ4 transcripts in CD4^+^ and CD8^+^ splenic T cells, albeit at low levels (34). In the context of osteoclasts, our microarray data revealed higher levels of PLCβ4 expression in mature osteoclasts than in their precursors. These results suggest the potential significance of PLCβ4 in osteoclastogenesis and bone homeostasis. Supporting this concept, *LysM-PLCβ4^−/−^* male mice displayed increased trabecular bone mass in their femurs and lumbar vertebra. The augmented bone mass observed in PLCβ4 deficiency was accompanied by a reduction in the number of osteoclasts and erosion area on the trabecular bone surface of the femurs, as measured by histomorphometry. Consistent with these *in vivo* findings, PLCβ4 deficiency led to reduced osteoclastogenesis *in vitro* when assessing BMMs with PLCβ4 knockdown by shRNA, as well as BMMs from *PLCβ4^−/−^* and *LysM-PLCβ4^−/−^* mice. On the other hand, the parameters representing osteoblast activity (MAR and BFR) did not exhibit significant changes in *LysM-PLCβ4^−/−^* mice. This suggests that PLCβ4 deficiency, specifically in osteoclast lineage cells, does not affect *in vivo* bone formation. Therefore, the increased bone mass observed in *LysM-PLCβ4^−/−^* mice was due to impaired osteoclastogenesis.

A previous study reported that *PLCβ4^−/−^* mice exhibited postnatal lethality due to motor defects (30). Consistent with our observations, *PLCβ4^−/−^* mice died prematurely at approximately 3 weeks of age. Furthermore, we found that the size of *PLCβ4^−/−^* mice at 8 weeks of age was smaller than that of WT mice. However, there were no significant differences in the skeletal development between the two groups at embryonic day 17.5, except for mandibular development, which indicates that PLCβ4 does not significantly influence bone development during the early stages. Additionally, when comparing body weights and size, no difference was observed between the WT and *LysM-PLCβ4^−/−^* mice at 8 weeks of age, which further confirms that the relatively small size of *PLCβ4^−/−^* mice was not because of defects in skeletal development and bone structure.

Interestingly, we observed sex-related differences in the bone phenotype in *LysM-PLCβ4^−/−^* mice. However, it is common to encounter sex-specific skeletal phenotypes in gene KO animal models, and similar results have been reported in previous studies (40–44). Our data unveiled that male mice exhibited alterations in their trabecular bone, whereas female mice did not show any changes. These findings demonstrate a sex-specific response in the absence of PLCβ4, although the precise underlying mechanism remains unclear. Further investigations are required to fully understand the reasons for the observed sex-specific effects on bone metabolism in the absence of PLCβ4.

Among the MAPKs, p38 plays a crucial role as an essential regulator of NFATc1 induction that is required for RANKL-mediated osteoclastogenesis, thus making it a potential target for treating osteoporosis. In this study, we observed that the phosphorylation of p38 in response to RANKL was stunted in the absence of PLCβ4. It is worth noting that PLCβ4 deficiency did not interfere with the activation of p38 MAPK induced by M-CSF or TNF-α, both of which are pivotal for osteoclast generation. These results demonstrate that PLCβ4 selectively facilitates RANKL-mediated p38 phosphorylation. This effect of PLCβ4 on the regulation of p38 activation emphasizes its critical role in mediating RANKL-induced osteoclast differentiation.

Phosphorylation of the NF-κB p65 subunit by RANKL is essential for NF-κB transcriptional activity (37, 45). Notably, treatment with a p38 inhibitor blocks p65 phosphorylation at Ser-536 by RANKL, which subsequently attenuates the transcriptional activity of NF-κB (37). These observations reveal that p38 MAPK is involved in modulating p65 phosphorylation in response to RANKL. Consistently, we observed a reduction in the phosphorylation of p65 at Ser-536 in *LysM-PLCβ4^−/−^* BMMs. This decrease in p38 and p65 phosphorylation correlated with the attenuated expression of osteoclastogenic markers, such as NFATc1, TRAP, and CTSK, in *LysM-PLCβ4^−/−^* cells. Therefore, the reduced phosphorylation of p38 and p65 by RANKL contributes to impaired osteoclastogenesis in the absence of PLCβ4. The function of p38 MAPK is regulated by two upstream MAPK kinases, MKK3 and MKK6, both of which are known to have an essential role in osteoclast differentiation. *In vivo* studies using *MKK3^−/−^* and *MKK6^−/−^* mice have shown that both groups exhibit increased bone mass because of impaired osteoclast formation (46). Interestingly, osteoclastogenesis was significantly decreased in the *MKK3^−/−^* group, whereas MKK6 deficiency had no *in vitro* impact on osteoclast formation. In addition, the level of p38 phosphorylation and expression of osteoclast marker genes, including NFATc1, were significantly attenuated in *MKK3^−/−^* cells but not in *MKK6^−/−^*cells. These findings indicate that MKK3, rather than MKK6, is a principal regulator of osteoclast differentiation *in vitro*. Our data revealed that PLCβ4 interacts with MKK3 and p38 to form a complex upon RANKL stimulation. In addition, no association between PLCβ4 and MKK6 was observed (data not shown). These data are consistent with those of a previous study using the yeast two-hybrid system and co-immunoprecipitation assays (38), which supports the observed interaction of PLCβ isozymes with MKK3 and p38 MAPK in this study. Immunofluorescence analysis further confirmed the colocalization of PLCβ4 and p38 following RANKL stimulation. Moreover, treatment with a p38 activator restored the defective osteoclast formation in the absence of PLCβ4. Interestingly, the PLCβ4 inhibitor U73122 did not influence p38 phosphorylation, thus providing evidence that the catalytic activity of PLCβ4 is not required for RANKL-mediated osteoclastogenesis. Collectively, these findings highlight the crucial role of PLCβ4 in assembling the components of the MKK3 and p38 MAPK signaling modules to induce p38 activation.

Our study provides convincing evidence that PLCβ4 plays a critical role in maintaining bone homeostasis through osteoclastogenesis regulation. We propose a working model whereby PLCβ4, functioning as an adaptor protein, mediates complex formation with MKK3-p38 and the activation of p38 and p65, thus facilitating osteoclast differentiation (Fig. 8d). Our findings emphasize the significance of PLCβ4 in osteoclast biology and suggest its potential as a therapeutic target for managing bone diseases that are characterized by excessive osteoclast formation.

## Materials and methods

### Mice

The generation of PLCβ4 global KO mice with a C57BL/6 background has been previously reported (30). Mice with disrupted PLCβ4 in LysM-expressing cells were generated by crossing LysM-Cre transgenic mice with PLCβ4-floxed mice, and both maintained their C57BL/6 background. All mice were kept in a specific pathogen-free mouse facility with access to sterilized food, water, and bedding under a 12-h/12-h light/dark cycle at 25°C. All animal procedures were reviewed and approved by the Committee on the Ethics of Animal Experiments of Kyungpook National University (Approval No. KNU-2019-0038). Animals were handled in accordance with the relevant guidelines.

### Reagents and antibodies

Recombinant human RANKL was purchased from R&D Systems (Minneapolis, MN, USA), and M-CSF was obtained from PeproTech (Rocky Hill, NJ, USA). The PLCβ4-specific antibody was purchased from Santa Cruz Biotechnology (Santa Cruz, CA, USA). Antibodies against phospho-IκBα, phospho-JNK, phospho-ERK, phospho-p38, phospho-p65, IκBα, JNK, ERK, p38, MKK3, and MKK6 were obtained from Cell Signaling Technology (Beverly, MA, USA). Antibodies specific for NFATc1 and cathepsin K were purchased from BD Pharmingen (San Diego, CA, USA) and Millipore (Billerica, MA, USA), respectively.

### Macrophage isolation and osteoclast culture

Osteoclasts were generated as previously described (47). Briefly, primary bone marrow-derived macrophages (BMMs) were isolated from the long bones of 8–9-week-old mice and cultured in α-minimal essential medium (MEM) supplemented with 10% fetal bovine serum in a culture dish. The following day, bone marrow stromal cells and red blood cells were removed, and only BMMs were collected and cultured in α-MEM containing 10% fetal bovine serum with 1/10 volume of CMG 14–12 cell culture medium as a source of M-CSF (48) in a Petri dish. After 3 days, the adherent cells were lifted and used as osteoclast precursors (BMMs). The BMMs were seeded at a density of 5×10^3^ cells/well in a 96-well cell culture plate and cultured with M-CSF (30 ng/mL) and RANKL (20 ng/mL) for 4–5 days to induce osteoclast formation. After the culture period, the osteoclasts were washed with phosphate-buffered saline (PBS) and fixed with 4% paraformaldehyde for 20 min. Subsequently, the cells were stained with TRAP solution, which consisted of 0.1M sodium acetate solution (pH 5.0) containing 6.67 mM sodium tartrate, 0.1 mg/mL naphthol AS-MX phosphate, and 0.5 mg/mL Fast Red Violet. TRAP-positive multinucleated cells containing three or more nuclei were counted as osteoclasts.

### Microarray analysis

BMMs were cultured for 4 days with 30 ng/mL of M-CSF and 20 ng/mL of RANKL. The resulting gene expression microarray data were analyzed using the Affymetrics gene expression software.

### Lentiviral transduction of PLCβ4 shRNA

The lentiviral constructs containing PLCβ4 small hairpin RNA (shRNA) or a non-specific shRNA control (Con-sh) were obtained from Sigma-Aldrich (St. Louis, MO, USA). Lentiviral particles were generated by transfecting 293T cells with the expression constructs, virus packaging plasmid (ΔH8.2), and envelop plasmid (VSVG) using FuGENE HD transfection reagent (Promega, Madison, WI, USA). The supernatant containing the constructed lentiviruses was collected after 24–48 h of transfection. BMMs were transduced with the viral supernatant along with 10 μg/mL of protamine sulfate (Sigma-Aldrich) for 24 h. Subsequently, the cells were subjected to selection in the presence of 4 μg/mL of puromycin for 3 days.

### Cell proliferation assay

Cell proliferation was determined using the 3-(4,5-dimethylthiazol-2-yl)-5-(3-carboxymethoxyphenyl)-2-(4-sulfophenyl)-2H-tetrazolium (MTS) assay. BMMs were cultured in 96-well plates at a density of 5 × 10^3^ cells/well with various concentrations of M-CSF. After 3 days, the existing media was removed and replaced with 100 µL of medium containing 20 µL of CellTiter 96 AQueous One solution (Promega). The cells were then incubated for 4 h at 37°C in a humified atmosphere with 5% CO_2_, and absorbance was measured at 490 nm using a microplate reader.

### Quantitative real-time polymerase chain reaction

Total mRNA was isolated from the cultured cells, and cDNA was synthesized as previously described (49). Quantitative real-time PCR was performed using an ABI 7500 real-time PCR system and SYBR Green dye (Applied Biosystems, Foster City, CA, USA). The mRNA expression levels were normalized to the endogenous GAPDH housekeeping gene. Primer sequences were as follows: PLCβ1, 5′-CTG CAC AGA GGA TGT GCT GA-3′ and 5′-CCA AGT GTC CGA TGT TCC CA-3′; PLCβ2, 5′-GCT TCC TCT CCT GTT CAC CC-3′ and 5′-CCT TCA CGT TAG GGG GCA AT-3′; PLCβ3, 5′-GGA GCG TGT GGA GAG AGC AG-3′ and 5′-AGC ACT TCG TTG AGT CTC GG-3′; PLCβ4, 5′-TGC CAG ATG GTT TCA CTG AA-3′ and 5′-GAA GGT ACC CGC ATG ATC CA-3′; NFATc1, 5′-ACC ACC TTT CCG CAA CCA-3′ and 5′-TTC CGT TTC CCG TTG CA-3′; TRAP, 5′-TCC CCA ATG CCC CAT TC-3′ and 5′-CGG TTC TGG CGA TCT CTT TG-3′; Cathepsin K, 5 ′-GGC TGT GGA GGC GGC TAT-3′ and 5′-AGA GTC AAT GCC TCC GTT CTG-3′; OSCAR, 5′-GCT GGC TGC GCT GTG AT-3′ and 5′-ACC TGG CAC CTA CTG TTG CT-3′; Atp6V0d2, 5′-GAG CTG TAC TTC AAT GTG GAC CAT-3′ and 5′-CTG GCT TTG CAT CCT CGA A-3′; DC-STAMP, 5′-CTT CCG TGG GCC AGA AGT T-3′ and 5′-AGG CCA GTG CTG ACT AGG ATG A-3′; and MITF, 5′-GCT ATG CTC ACT CTT AAC TCC AAC-3′ and 5′-TTG GGG ATC AGA GTA CCT AGC TCC-3′

### Immunoblotting and immunoprecipitation

Cultured cells were washed with ice-cold PBS and lysed in lysis buffer [50 mM Tirs-HCl (pH 7.4), 150 mM NaCl, 1% Nonidet P-40, 1 mM EDTA] containing protease and phosphatase inhibitors for 10 min on ice. The cell lysates were centrifuged at 13,000 rpm for 30 min at 4°C. Protein concentrations of the cell lysates were measured using a Bicinchoninic Acid Kit (Thermo Scientific Inc., Rockford, IL, USA). An equal amount of quantified protein for each sample was subjected to 8% or 10% SDS-PAGE and transferred to polyvinylidene difluoride (PVDF) membranes. After blocking with 5% skim milk or 3% bovine serum albumin (BSA), the membranes were incubated overnight with the indicated primary antibodies at 4°C, followed by probing with the appropriate secondary antibodies. Blots were visualized using an ECL-Plus detection kit (Amersham Pharmacia Biotech, Piscataway, NJ, USA). For immunoprecipitation, the cell lysates were incubated with an anti-PLCβ4 antibody or control IgG, followed by Sepharose A beads (GE Healthcare). The immunoprecipitated proteins were separated by 10% SDS-PAGE and immunoblotted as described above.

### Microcomputed tomography (µCT)

Mouse femurs were isolated and fixed in 4% paraformaldehyde for 24 h. Subsequently, the femurs were scanned using a Quantum FX μ-CT (Perkin Elmer, Waltham, MA, USA) with a voxel resolution of 9.7 μm, operating at 90 kV and 200 μA. The field of view was set at 10 mm, and the exposure time was 3 min. The region of interest was defined as 0.3 mm from the bottom of the growth plate. Three-dimensional models were constructed from the obtained scan data, and the bone parameters were determined using Analyze 12.0 software (Overland Park, KS, USA).

### Histology and bone histomorphometric analysis

For histological analyses, the proximal tibiae were isolated and fixed in 4% paraformaldehyde for 24 h to evaluate the *in vivo* osteoclast parameters. Subsequently, decalcification was performed in 10% EDTA for 4 weeks at 4°C. Following decalcification, the bones were dehydrated, embedded in paraffin, and sectioned. The sections were stained with TRAP to visualize the osteoclasts. For dynamic bone histomorphometry analysis, the mice were administered intraperitoneal injections of calcein green (10 mg/kg) and alizarin red (20 mg/kg) on the sixth and second days before being sacrificed. The lumbar vertebrae were fixed and embedded in methyl methacrylate resin without decalcification (47). The resin blocks were cut longitudinally into 6-μm slices of the tibia using a Leica RM2165 rotary microtome equipped with a tungsten blade (Leica Microsystems, Germany). Fluorescence signals emitted by calcein green and alizarin red were recorded using a fluorescence microscope (Leica Microsystems). For von Kossa staining analysis, the lumbar vertebrae were fixed and embedded in methyl methacrylate. Subsequently, 6-μm-thick samples were stained with von Kossa reagent. All quantitative histological parameters were evaluated using the Bioquant OSTEO II program (Bio-Quant, Inc., Nashville, TN).

### Immunofluorescence assay

Cells cultured on glass slides in a 24-well plate were fixed with 4% paraformaldehyde for 20 min, permeabilized with 0.1% Triton X-100, and blocked with 0.2% BSA for 10 min. The cells were then incubated for 2 h with the primary antibodies in a solution containing 0.2% BSA. To visualize the binding of primary antibodies, fluorescent dye-conjugated secondary antibodies (Molecular Probes, Eugene, OR, USA) were applied in a 0.2% BSA solution and incubated for 1 h. Immunofluorescence-labeled cells were observed using a fluorescence microscope (Leica Microsystems).

### Statistical analysis

Statistical analyses were performed using Microsoft Excel 2016 (Microsoft, USA) or GraphPad Prism 9 (GraphPad, San Diego, CA, USA). Statistically significant differences between the two groups were determined using the Student’s *t*-test. The data are presented as the mean ± standard deviation (SD) or mean ± standard error (SE). A *p*-value of < 0.05 was considered statistically significant.

## Acknowledgments

This work was supported by the National Research Foundation of Korea (NRF) grant funded by the Korean Government (MSIT) (NRF-2018R1A2B6001298 to H.-J.K. and NRF-2022R1A2C1006105 to J.-Y.C.) and the Basic Science Research Program through the NRF funded by the Ministry of Education (NRF-2021R1I1A1A01051983 to H.-J.K.).

