## Supplementary material for "Phospholipase C β4 promotes RANKL-dependent osteoclastogenesis by interacting with MKK3 and p38 MAPK": Fig. S1

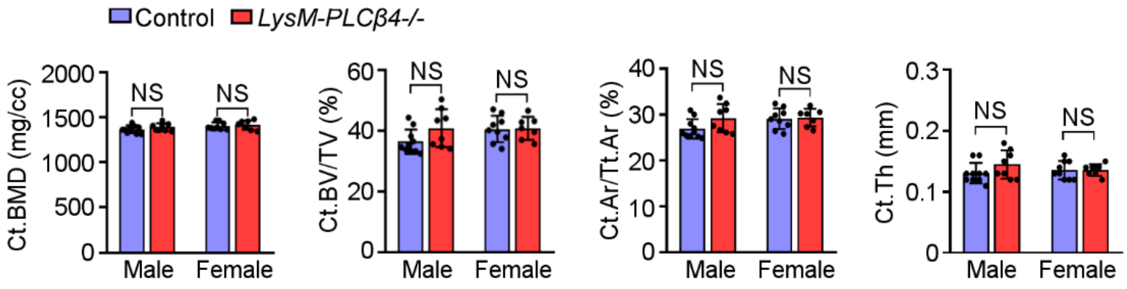


**Fig. S1** Deletion of *PLCβ4* in the osteoclast lineage does not affect cortical bone parameters. Quantitative μCT analysis of cortical bone parameters of femurs from 8-week-old control and *LysM-PLCβ4*$^{-/-}$ male and female mice. Quantitative measurements of cortical bone mineral density (Ct.BMD), cortical bone volume per tissue volume (Ct.BV/TV), cortical area per total area (Ct.Ar/Tt.Ar), and cortical thickness (Ct.Th) (n = 7–10). NS, not significant
